# MAVeRiC-AD: Mixture-of-experts Agentic Vision-Language Ensemble for Robust MRI Classification of Alzheimer’s Disease

**DOI:** 10.1101/2025.09.05.674598

**Authors:** Nikhil J. Dhinagar, Pavithra Senthilkumar, Sophia I. Thomopoulos, Paul M. Thompson

## Abstract

Robust classification of Alzheimer’s disease (AD) from structural T1-weighted MRI (T1w) images remains an unmet clinical need, especially when data is acquired at multiple sites that differ in scanning protocols and population demographics. In this paper, we present MAVeRiC-AD (Mixture-of-experts Agent-guided Vision-Language Ensemble for Robust Imaging-based classification of Alzheimer’s Disease), an agentic framework that dynamically utilizes the optimal inferencing tool for radiological queries to provide relevant answers to the user. In our framework, we tested three specialized models encoded as callable tools: (1) CNN-AD, a 3D DenseNet trained on T1w intensities only; (2) MOE-VLM, a vision-language model that jointly models the T1w with subject-specific demographics (age, sex, site) via a mixture-of-experts (MoE) projection head; (3) Retrieval engine, a similarity-search module that contextualizes a patient against others from the site and reports % prevalence of AD. A light-weight agent analyzes the user request and then routes the input (image, or image + text) to the appropriate tool, aggregates responses and returns the tool response augmented with its confidence derived from conformal prediction. Experiments were conducted using T1w images from the ADNI (N=4,098) and OASIS-3 (N=600) datasets. Single-site training baselines achieved ROC-AUC = 0.79 (CNN) and 0.82 (VLM) on ADNI. When trained jointly on both sites, MOE-VLM surpassed both image-only and standard vision-language models with ROC-AUC = 0.90 on ADNI and 0.81 on OASIS. MAVeRiC-AD demonstrates that agentic orchestration of complementary expert deep models, coupled with explicit demographic conditioning for multi-site data can improve robustness and interpretability of AD image analysis pipelines and serves as a blueprint for scalable, trustworthy clinical AI assistants.

## 1. DESCRIPTION OF PURPOSE

Alzheimer’s disease (AD) affects over 55 million people worldwide; robust and reliable diagnosis is critical for patient management. Magnetic-resonance imaging (MRI) is a non-invasive biomarker, yet automated classifiers trained on single-site data often fail to generalize across institutions because of scanner variability and demographic shifts. Recent vision-language models (VLMs) align images with natural language-based descriptions, offering a principled way to use demographic context [1]. Simultaneously, clinical AI systems are moving towards agentic paradigms—interactive assistants capable of calling specialized tools, explaining their reasoning along with their answers. An “ agent,” is usually defined as a system that iteratively reasons– acts-observes it reads inputs and intermediate results, reasons internally, and then invokes relevant expert tools. ReAct-style agents [2] interleave natural-language reasoning with tool calls, leaving an auditable trail of decisions and enabling verification of each step. In clinical workflows, such agents can validate inputs, ask clarifying questions, and route a case to the model with the most relevant expertise. Frameworks like VoxelPrompt [3], AD-Agent [4] and VILA-M3[5] underline this trend, yet recent works either operate on 2D data, or ignore demographic priors.

A limiting factor for cross-site robustness is the use of dense projection layers that treat all inputs identically. Mixture-of-Experts (MoE) layers replace a single feed-forward block with *E* parallel “ experts,” each a small neural network. A gate network usually learns a soft-max routing function and forwards tokens to the *k* experts with highest weights, enabling specialization (e.g., sex-specific or site-specific experts) while keeping computation sparse. Some of the original pioneering work of MoE models in practice was by Hinton et al. [6] and Sutskever et al. [7]. Recent flagship foundation models for language and vision such as OpenAI’s gpt-oss [8], Llama 4 [9], Qwen 3 [10], and Kimi-VL [11] utilize an MoE style architecture.

We therefore introduce MAVeRiC-AD—a single ReAct agent that:

1. **Parses user intent** (classification, relative site AD prevalence) and **available context** (image, age, sex, site).
2. **Routes** to one of three tools: an image-only CNN, a **demographic-aware** VLM, or our **MoE-VLM** with top-*k* gating.

Across ADNI & OASIS-3, the MoE-VLM outperforms both CNN and standard VLM (AUC ≈ 0.90 vs 0.79– 0.88 and 0.81 vs 0.71-0.79) while providing interpretable expert-utilization visualizations. MAVeRiC-AD thus offers a practical path toward trustworthy, transparent, multi-tool neuro-AI assistants.

**Figure 1.**
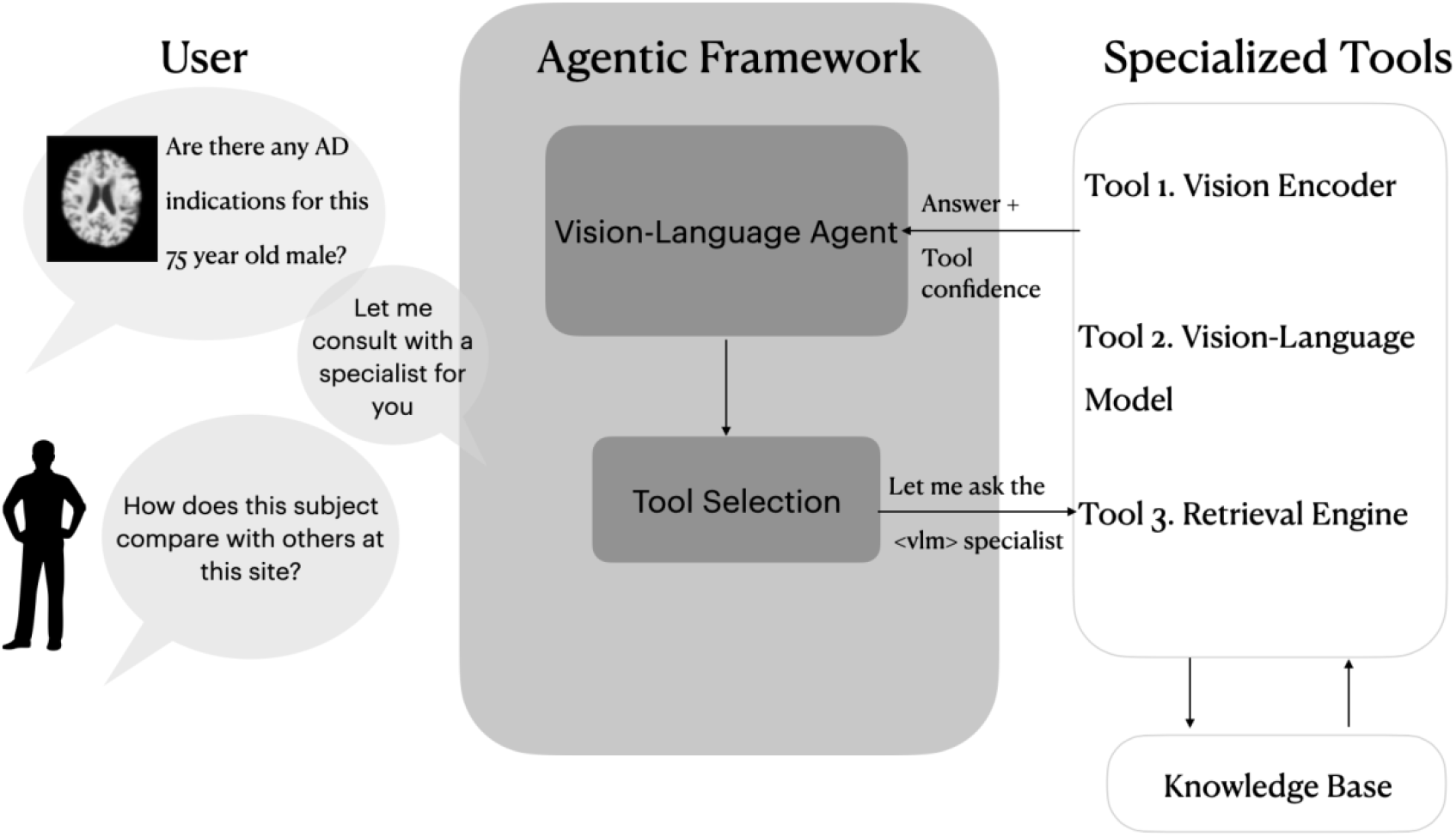
MAVeRiC-AD, the proposed single agentic framework that can take image-only, or image + text as input, and automatically choose the best analysis pathway.

## 2. METHODS

### 2.1. Problem Definition

We study binary diagnosis of Alzheimer’s disease (AD) versus cognitively normal (CN) controls from T1w MRI acquired across multiple sites. Each subject has a 3D T1-weighted MRI volume and optional demographic/site metadata (age, sex, acquisition site) that are used as inputs. For vision–language models (VLMs), metadata are formatted as a text prompt (e.g., “ 75-year-old male subject diagnosed with Alzheimer’s from site ADNI”). The primary objective for the framework is to classify each test subject, i.e., p(y=AD|image, text) and return its confidence based on conformal prediction. Key challenges include (i) the distribution shift across sites and scanners, (ii) limited per-site dataset sizes, and (iii) the need for user transparency.

### 2.2 Vision-language Agent

The agent mediates interactions between the user and the underlying models/tools.

#### Request parsing

The agent identifies the user goal (classification, relative site AD prevalence) and extracts data (image, and availability of age, sex, site).

#### Routing

The agent is based on the TinyLlama 1.1B LLM [12] (based on the architecture and tokenizer in Llama2) finetuned using low-rank adaptation (LoRA) on N=600 synthetic prompt–tool pairs (generated by prompting the Mistral 7B LLM [13]) that map semantic variations of user instructions (image ± age/sex/site) to tool tags (<cnn>, <vlm>, <retrieval>).

- If only an image is provided → route to the image-only CNN.
- If image and demographics/site are provided → route to the VLM (standard VLM or MoE-VLM).
- If the user requests cohort context (e.g., “ Given this 75 year old male subject’s T1w, retrieve a summary of similar subjects and the AD prevalence in ADNI?”) → route to the retrieval tool.

#### ReAct orchestration with LangGraph

The agent is implemented as a LangGraph state graph following the **ReAct** pattern: the control flow alternates between *Thought (llm) → Action (tool call) → Observation (back to llm)*. The state is the running memory for the agent to keep track of tool calls and tool outputs. Graph nodes are functions that help move between states i.e., *Invoke Tool* (CNN/VLM/Retrieval). Edges are chosen from observations, to decide which node to call next without hand-coding the flow.

#### Orchestration and feedback

The agent standardizes inputs (preprocessing, prompt construction), calls the selected tool, and returns a structured report: predicted class, probability, conformal certainty (“ Certain/Ambiguous”), and optional site AD prevalence based on the input subject information. Errors (e.g., missing files and unclear data formats) trigger clarifying prompts rather than silent failures.

### 2.3 Individual Tools: image only, image + text, retrieval

#### Image-only CNN

Here a trained 3D convolutional network (initialized from supervised pretraining) outputs p(AD| image). It served as a baseline and a fallback for the agent when the text-based demographics were not provided with the T1w image.

#### Standard VLM (image+text)

This VLM setup follows the methodology described in this recent paper [1]. A dual encoder mapped the MRI and text prompt to a joint embedding space; The model learns this optimal space by maximizing the normalized cosine similarity between the correct pair of T1w image and corresponding text prompt. The text prompts were augmented at training time with semantically equivalent phrasings to reduce prompt overfitting.

#### MoE-VLM (image+text)

We replaced each projection head with a Mixture-of-Experts (MoE) style head where a gate network routed the tokens through *k* parallel experts. The gate took as input image/text embeddings from the specialized encoders and produced routing weights. We used top-k routing during training where the sparsity encouraged specialization (age, sex, diagnosis, and site-specific) without manual hand-coding of features.

#### Retrieval

Given the test subject’s T1w image and additional provided context (age, sex, site), the tool summarizes AD-prevalence as percent in the site based on a similarity-based retrieval using the image and filtered by the context.

## 3. EXPERIMENTAL SETUP

### 3.1 Data

In line with similar studies [14], all 3D T1-weighted brain MRI scans were pre-processed here using standard steps for neuroimage analyses including nonparametric intensity normalization (N4 bias field correction) [15], ‘skull-stripping’, linear registration to a template with 9 degrees of freedom, and isotropic voxel resampling 2-mm resolution. All images were *z*-transformed (setting each image’s mean and SD to a standard value) to stabilize model training. The T1-w images were re-sized to 80 voxels across all dimensions before model training. We used data from ADNI [16] and OASIS-3 [17] datasets as the two unique imaging sites for each of our experiments for AD classification. The datasets are summarized in **Table 1** below.

**Table 1.** Summary of data utilized in this work.

| Dataset | N | #Subjects | Age (years) | Sex (F/M) |
| --- | --- | --- | --- | --- |
| ADNI | 4,098 | 1,188 | 55.7-92.8 | 556F/632M |
| OASIS | 600 | 600 | 43.5-97.0 | 341F/359M |

### 3.2 Model Training and Testing

We performed a random search to select hyperparameter values. The embedding dimension was 1024 for the 3D CNN based [18] vision encoder and 384 for the sentence BERT [19] based text encoder. The models were trained with a batch size of 64, AdamW optimizer. All models minimized the binary cross entropy loss functions for classification. We evaluated our models with the receiver-operator characteristic curve-area under the curve (ROC-AUC).

## 4. RESULTS

Table 2 summarizes results of specialist tools and related ablation experiments. Additionally, we tested the capability of the agent to invoke each tool with simulated user prompts as well as paraphrased versions of the prompts to test generalizability. In each of our test experiments the agent was able to correctly route the user request to the appropriate tool.

## 5. DISCUSSION

Our results in **Table 2** show clear benefits from combining vision–language modeling with a mixture-of-experts style architecture and the capability of the agents to leverage tools based on them. The MOE-VLM with prompt augmentation and top-k routing achieved the strongest ADNI performance (AUC = 0.902) and the best cross-site result on OASIS (AUC = 0.812), outperforming the image-only CNN (0.790/0.709) and standard VLM model. The performance gains likely arise from (i) the natural language supervision to provide additional context and (ii) expert specialization using k-gating in the MoE style architecture. The improvement from text prompt augmentation suggests text diversity reduces overfitting to a single template. The agent was able to accurately invoke the appropriate tools based on the user prompt in our experiments.

**Table 2.** Summary of the specialist tools (image only, image +text) performance for AD classification. The Mixture-of-experts (MOE) VLM outperformed the standard VLM on multi-site data.

|  | Training | CNN pretrain | Test ADNI<br>(N = 1219) | Test OASIS<br>(N = 60) |
| --- | --- | --- | --- | --- |
| **MOE-VLM + image and prompt augment, k-gate** | Joint | ADNI super | 0.902 | 0.812 |
| **MOE-VLM + image and prompt augment** | Joint | ADNI super | 0.903 | 0.787 |
| **VLM + image and text prompt augment,** | Joint | ADNI super | 0.902 | 0.796 |
| **VLM** | Joint | ADNI super | 0.878 | 0.802 |
| **VLM [1]** | ADNI | ADNI super | 0.821 | - |
| **CNN (image only)** | Joint | none | 0.790 | 0.709 |

## 6. CONCLUSION

In this paper, we proposed MAVeRiC-AD, an agentic framework that parses the user request in natural language and then autonomously orchestrates the relevant use of multiple specialized deep neural networks for Alzheimer’s disease classification and retrieval. We tested and integrated a MoE head inside the vision-language model to make use of the available context (age, sex, site) in addition to the T1w image. The specialized tools in our framework yield competitive multi-site AD classification results while automatically flagging uncertain cases. MAVeRiC-AD illustrates how agentic tool selection, and expert specialization together advances trustworthy medical-imaging AI.

